# Sexual morph specialisation in a trioecious nematode balances opposing selective forces

**DOI:** 10.1101/2021.08.31.458370

**Authors:** Sally Adams, Prachi Pathak, Maike Kittelman, Alun R.C. Jones, Eamonn B. Mallon, Andre Pires-daSilva

## Abstract

The coexistence of different mating strategies, whereby a species can reproduce both by selfing and outcrossing, is an evolutionary enigma that has long intrigued biologists (Darwin, 1877). Theory predicts only two stable mating states : outcrossing with strong inbreeding depression or selfing with weak inbreeding depression. As these two mating strategies are subject to opposing selective forces, mixed breeding systems are thought to be a rare transitory state, yet they have been found to persist even after multiple speciation events. We hypothesise that if each mating strategy plays a distinctive role during the species life history, opposing selective pressures could be balanced, permitting the stable co-existence of selfing and outcrossing sexual morphs. In this scenario, we would expect each sexual morph to be specialised in their respective roles. Here we show, using a combination of behavioural, physiological and gene expression studies, that the selfing (hermaphrodite) and outcrossing (female) sexual morphs of the trioecious nematode *Auanema freiburgensis* have distinct adaptations optimised for their different roles during the life cycle. *A. freiburgensis* hermaphrodites are produced under stressful conditions, are specialised for dispersal to new habitat patches and exhibit metabolic and intestinal changes that enable them to meet the energetic cost of dispersal and reproduction. In contrast, *A. freiburgensis* females are produced in favourable conditions, facilitate rapid population growth and compensate for the lack of reproductive assurance by reallocating resources from intestinal development to robust mate-finding behaviour. The specialisation of each mating system for their role in the life cycle could balance opposing selective forces allowing the stable maintenance of both outcrossing and selfing mating systems in *A. freiburgensis*.

## INTRODUCTION

Natural selection has driven the evolution of diverse modes of reproduction, ranging from species that replicate exclusively from a single parent to those that have separate sexes. The coexistence of different mating strategies within a species, where conspecifics can reproduce either by outcrossing (male and female) or self-fertilising (hermaphrodites), is an evolutionary enigma that has long intrigued biologists <u>(Darwin,1877)</u>. Species with such mixed mating strategies are usually considered a temporary transitional state, as theory predicts only two stable mating states: outcrossing with strong inbreeding depression or selfing with weak inbreeding depression (Charlesworth, 1984; Charlesworth et al., 1990; Lande and Schemske, 1985). Yet mixed breeding have been found to persist even after multiple speciation events (Weeks, 2006), suggesting that in some cases they are not transitory.

Mixed mating strategies are expected to be rare, as they are subject to opposing selective forces. In a transitory system, self-fertilising hermaphrodites should outcompete females and males if there is strong selection for reproductive assurance (guaranteeing reproduction even if a male is not available) (Pannell, 1997a; Wolf & Takebayashi, 2004). Otherwise, dioecy (females and males) should dominate if selection for reproductive assurance is relaxed, e.g. to reduce inbreeding or in response to selection for sexual morph specialization, reviewed in (Charlesworth, 1999). We would expect a stable mixed mating system to exist only where these opposing selective forces are balanced.

To date, the study of sexual polymorphisms has primarily focused on flowering plants. However, the large and diverse phylum Nematoda provides an opportunity to study the evolution and potential stability of mixed mating strategies and its relationship to specific life histories. Although dioecy is the most common mode of reproduction in nematodes, several other mating systems have evolved, including hermaphroditism, androdioecy (males and hermaphrodites) and trioecy (males, females and hermaphrodites) (Denver et al., 2011; Pires-daSilva, 2007). The recently described nematode genus *Auanema* (Kanzaki et al., 2017) displays trioecy, having two egg-laying reproductive modes that coexist in a population: females, which must outcross with males to reproduce, and hermaphrodites, which although able to cross with males, predominantly reproduce by self-fertilisation (Tandonnet et al., 2018). In *A. freiburgensis*, sex determination between non-males (females and hermaphrodites) and males is regulated chromosomally: males inherit a single X chromosome (XO), whilst hermaphrodites and females inherit two (XX) (Figure 1). *Auanema* males are under-represented in populations, as they are produced at low percentages both from male-female crosses (< 20%) and self-fertilising hermaphrodites (< 10%) (Chaudhuri et al., 2015; Félix, 2004; Kanzaki et al., 2017; Shakes et al., 2011; Winter et al., 2017). In contrast, environmental cues play an important role in determining the non-male sexual morphs in *A. freiburgensis* (Robles et al., 2021; Zuco et al., 2018). Whereas at low population density hermaphrodites produce mostly female progeny, sensing of a crowding signal triggers the hermaphrodite mother to switch to produce predominantly hermaphrodite offspring. The female and hermaphrodite exhibit different larval development (Figure 1). When in well-fed conditions at 20 °C, a hermaphrodite takes approximately 3 days from hatching to adulthood. The hermaphrodite obligatorily passes through the dauer stage, a specialised non-feeding, stress-resistant, migratory larva. In contrast, the *A. freiburgensis* female and male never pass through dauer and reach adulthood after approximately 2 days at 20 °C.

**Figure 1.**
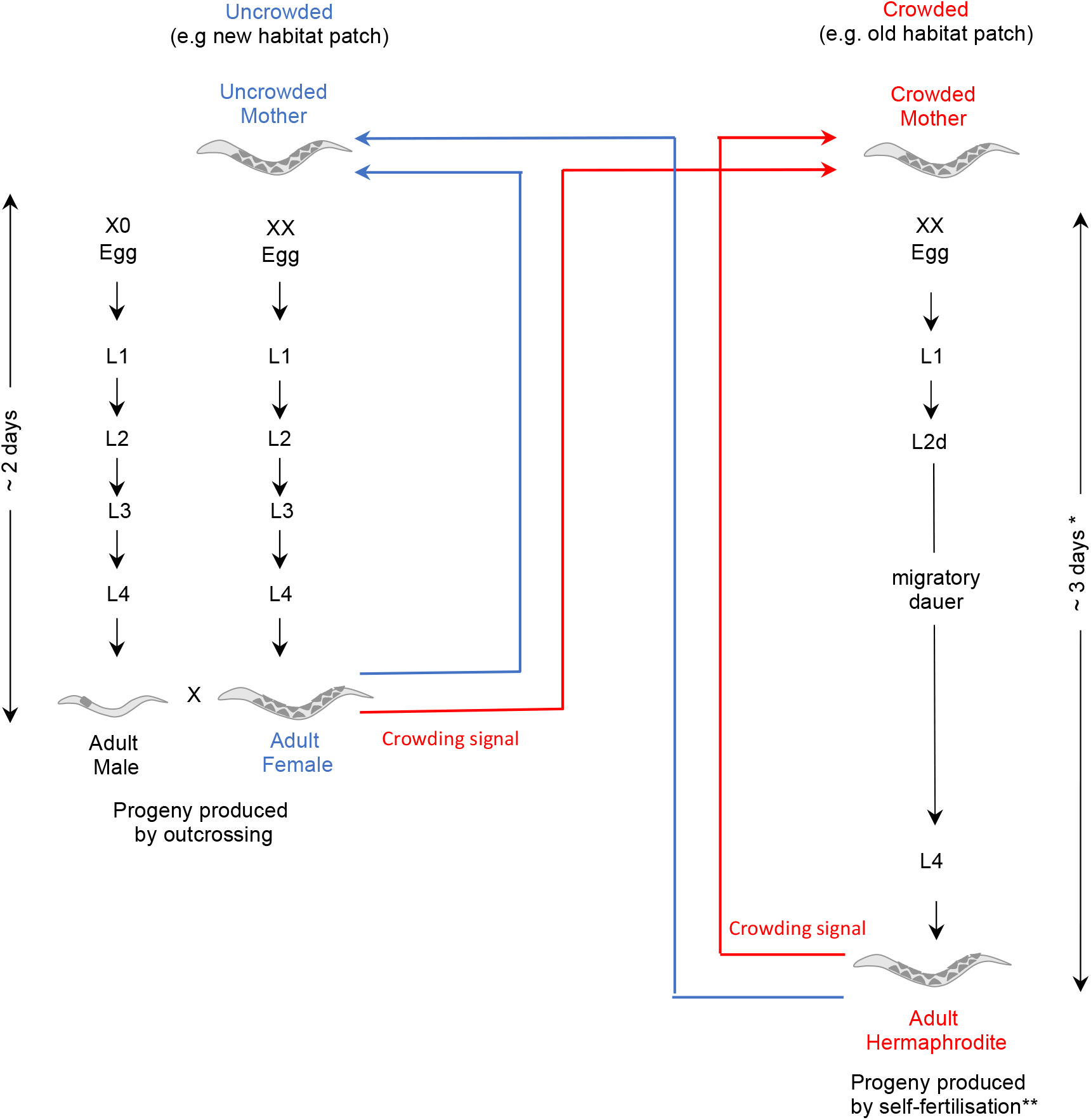
The life cycle of *A. freiburgensis*. In *A. freiburgensis*, whether a mother produces females or hermaphrodite progeny depends upon her perception of her habitat. In uncrowded conditions (blue arrows), she produces outcrossing female progeny, but upon sensing a crowding signal can switch to producing selfing hermaphrodites (red arrows). Hermaphrodite progeny pass through an obligatory dauer larval stage optimised for migration. If dispersed to a new habitat patch the dauer will complete reproductive development and produce progeny by self-fertilisation. If this new habitat patch is uncrowded the hermaphrodite will produce mostly female (and male) progeny (blue arrows). As neither females or males pass through the dauer diapause larval stage they reach sexual maturity quicker, which could allow more rapid population growth in new habitats. Times for development are based on well fed conditions at 20°C. *The dauer diapause can persist for longer in sub-optimal conditions. \*\**A. freiburgensis* hermaphrodites predominantly reproduce by self-fertilisation although they are able to outcross with males. Hermaphrodites are not able to cross with females.

The passage through dauer of the hermaphrodite is probably an adaptation to colonise new environments, assuring the establishment of a new population out of a single self-fertilising individual. *A. freiburgensis* hermaphrodite founders, when in low population density, will produce predominantly male and female progeny. Since females and males do not pass through dauer, subsequent population growth by the offspring of this founding generation will be more rapid. Outcrossing between the progeny of co-founders could facilitate adaptation by generating genetic variation (Colegrave et al., 2002; Goddard et al., 2005; Gray and Goddard, 2012; Poon and Chao, 2004).

We hypothesise that the female (outcrossing) and hermaphrodite (predominantly selfing) sexual morphs of *A. freiburgensis* play distinct roles during the life cycle and that each will therefore exhibit adaptations to minimise the cost of their specific life history. This would suggest that selection for reproductive assurance takes place during the initial colonisation phase (hermaphrodite) and selection for sex specialisation in subsequent generations (females). The temporal balancing of these opposing selection forces would allow the stable maintenance of trioecy. If this were the case, we would expect to see hermaphrodites and females specialise in their respective roles. To test our hypothesis, we will search for a series of behavioural, physiological and gene expression specialisations between females and hermaphrodites.

## MATERIALS AND METHODS

### Nematodes strains and culture

For all experiments, we used the inbred strain APS7 of *A. freiburgensis* (Kanzaki et al., 2017). This was maintained on nematode growth medium (NGM) agar plates, seeded with the streptomycin-resistant *Escherichia coli* OP50-1 strain and cultured at 20 °C, using standard *C. elegans* protocols (Stiernagle, 2006). Microbial contamination was prevented by adding 200 µg/mL nystatin and 100 µg/mL streptomycin to the NGM (Avery, 1993).

### Isolation of the different stages and sexual morphs of *A. freiburgensis*

*A. freiburgensis* dauer larvae invariantly develop into hermaphrodites (Kanzaki et al., 2017). Thus, to isolate hermaphrodites, dauers were identified on well-populated plates by their distinctive morphology (Kanzaki et al., 2017) and incubated on NGM OP50-1 plates until reaching adulthood (∼48 h after collection).

*A. freiburgensis* hermaphrodites, in uncrowded conditions, produce mostly female and male progeny (Kanzaki et al., 2017). To collect females, young adult hermaphrodites were allowed to lay eggs at a low population density (3 hermaphrodites per 2 cm diameter OP50-1 bacterial lawn). 24 to 36 h after egg laying began, L2 stage female larvae were distinguished from males by their different tail morphology and transferred to female-only plates to prevent fertilisation. From these plates, L4 or early adult females could be identified after ∼16 h or ∼24 h respectively.

To generate mated females (MF) L4/early adult females were co-cultured with males overnight (16 h). Crosses were carried out in a ratio of 5 females to 1 male (total 30 individuals per plate). This ratio was chosen to reflect that males only constitute ∼20 % of the *A. freiburgensis* population (Kanzaki et al., 2017) and to minimise male induced damage to females.

### Behavioural assays

#### Chemotaxis assay

Chemotaxis assays were performed as described previously (Chaudhuri et al., 2015). In brief, conditioned medium was generated by placing 20 young adult males in 100 μl of M9 buffer (Stiernagle, 2006) for 16 h at 20 °C. Samples were centrifuged at 15 000 rpm for 5 min to pellet the nematodes and the supernatant removed and used for the assay. Assays were conducted on NGM plates without bacteria. A spot of 5 μl of conditioned media and one of 5 μl of M9 buffer were added 3 cm apart, with the midpoint as the centre of the plate. Adult nematodes, at the described stage of development, were placed at the midpoint and scored for their location after 60 min. A total of 65 replicates were carried out, with a minimum of five replicates carried out with each sexual morph. To calculate the chemotaxis index (CI) the number of nematodes attracted to the test spot was subtracted by the number of nematodes in the control spot, and divided by the total number of nematodes assayed (Bargmann and Horvitz, 1991).

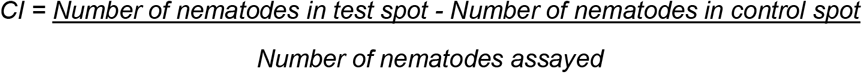

As the data was zero-inflated (animals not attracted to the male pheromone), we analysed it in two parts. We analysed if the animals were attracted to the male pheromone (yes/no) as a logistic regression, we then analysed how attracted they were using a generalized linear model with a gamma distribution in R. *Post-hoc* tests were carried out using the emmeans package version 1.5.4 in R (Lenth, 2021).

#### Leaving assay

Each sexual morph was isolated as described above. On the first day of adulthood, all individuals were moved to a fresh bacterial lawn (1 cm diameter in a 6 cm Petri dish) with 30 individuals per plate. The number remaining on the bacterial lawn was counted, 24 h and 48 h after the move, and expressed as a percentage of the starting number of individuals. This was replicated five times for each day and sexual morph combination, except day1:hermaphrodite which had three replicates (28 replicates in total). This data was analysed with a beta regression model in R (Cribari-Neto and Zeileis, 2010). Post-hoc tests were carried out using the emmeans package in R (Lenth, 2021).

### Physiological assays

#### Measurement and analysis of intestinal development

For all measurements individuals were imaged with the Zeiss Axio Zoom V16 (Zeiss) using the Zen 2 software (v2). Images were then converted to TIFF format and analysed in ImageJ. The *A. freiburgensis* intestinal cells are highly pigmented and easily distinguished. To determine the intestine width to nematode width ratio, 80 individuals were measured for both body width and intestinal width at the midpoint of third intestinal cell (from the posterior). This cell was easily identified in all stages and morphs. To determine the lumen length to nematode length ratio, 73 individuals were measured. The intestinal lumen is characterised by the lack of pigmentation and runs as a central line along the individual. Nematode length was measured from mouth to tail. All three measurements were analysed with linear models with a Gaussian distribution in R. We carried out Tukey’s posthoc honestly significant difference (HSD) tests in R.

#### Transmission electron microscopy (TEM)

Approximately 25 nematodes were placed into 100 μm aluminium platelets filled with OP50-1 *E. coli*. They were then frozen in a HPM-010 High Pressure Freezing machine (BalTec). Samples were processed in an automatic freeze substitution machine (RMC) in the following conditions: 90 °C for 14 h in 0.1% tannic acid in acetone, 72 h in 2 % OsO4 in acetone while slowly increasing temperature to 4 °C. Nematodes were further stained with 0.1 % thiocarbohydrazide in acetone for 2 h at room temperature, 2 h in 1 % OsO4 and infiltrated with increasing concentrations of 812 hard resin (Taab) over 4 days. Nematodes were finally thin-embedded between glass slides in 812 hard resin and polymerised at 70 °C for 24 hrs. 50 nm sections were cut with a PowerTome ultramicrotome (RMC) and examined with a Hitachi H-7650 transmission electron microscope operating at 100 kV. Microvilli were measured through the middle of the microvilli from the level of the apical base to the tip of the microvilli. At least 5 microvilli were measured per individual and the mean length recorded. In total, images from 27 (13 virgin female and 14 hermaphrodite) individuals were analysed. Unfortunately, mated females and later developmental stages could not be analysed due to sample deterioration.

#### Nile red staining and quantification

To measure neutral lipid stores we used the fixed Nile red staining method modified from the protocol by (Pino et al., 2013). Nile red (MP BIOMEDICALS SAS) was prepared as a 0.5 mg/ml stock in acetone. Staining solution was prepared just before use by adding 6 μl of Nile red stock solution per 1 ml of 40 % isopropanol (3 μg/ml final concentration). Approximately 50 nematodes per sample were collected in 300 μl of M9 buffer (Stiernagle, 2006). Samples were briefly washed in M9 buffer, to remove contaminating bacteria, and then fixed in 300 μl of 40 % isopropanol for 5 mins at room temperature. The isopropanol was removed and replaced with 300 μl of the Nile red staining solution. Samples were wrapped in foil and left at room temperature for 1 h. After staining, samples were washed twice with M9 buffer and mounted on microscope slides on agarose pads (2 % v/v agarose). Neutral lipid stores were visualised with the GFP filter set (excitation 482, emission 505) using the Zeiss Axio Zoom V16 (Zeiss). Images were taken at the specified exposure times and processed using the Zen 2 software (v2). To quantify corrected total nematode fluorescence (CTNF) for 146 individuals (mean of eighteen per possible sexual morph : day combination), images were converted to TIFF and processed with ImageJ. The CTNF was calculated per individual as described in (Hakim et al., 2018).

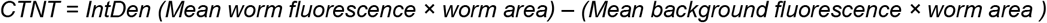

This data was analysed with linear models with a Gaussian distribution in R. We carried out Tukey’s posthoc HSD tests in R.

#### The chemical manipulation of dauer development

In nematodes, entry into dauer is regulated by the binding of steroidal endocrine hormone Δ7-dafachronic acid (DA) to the nuclear hormone transcription factor DAF-12 (Motola et al., 2006; Ogawa et al., 2009; Wang et al., 2009). In normal development DAF-9 (a cytochrome P450) synthesises high levels of DA, thereby blocking entry into dauer. In dauer-inducing conditions, DAF-9 inhibition results in low DA levels, facilitating DAF-12 mediated promotion of dauer development (Hu, Patrick, 2007). Addition of endogenous DA blocks dauer (Chaudhuri et al., 2011) while addition of specific inhibitor of DAF-9 (dafadine-A) forces entry into dauer (Luciani et al., 2011).

### Δ7-dafachronic acid (DA) inhibition of dauer development

DA-mediated inhibition of dauer development was carried out as described in (Chaudhuri et al., 2011). (25S)-Δ7-Dafachronic Acid (Cambridge Bioscience Ltd) was prepared as a 1 mM stock in 100 % ethanol. Immediately before use, the DA stock was diluted in water to 10 µM DA (1% ethanol). 20 µl of 10 µM DA or 1% ethanol alone (control) were added directly to a 1 cm diameter *E. coli* OP50-1 lawn on NGM plates and plates dried for 2 h. Under crowded conditions, *A. freiburgensis* hermaphrodite mothers predominantly produce hermaphrodite offspring (Kanzaki et al., 2017). Therefore, eggs were transferred from crowded plates, to either DA or control plates (4 plates for each condition with approximately 30 eggs per plate) and incubated at 20 °C for 48 h, before the sex of the progeny was determined by phenotype. All non-male progeny was subsequently moved to NGM plates (without DA or ethanol) to continue development and examined daily for intestinal phenotype. In total, 630 individuals were scored (409 with DA, 221 control) from 3 individual experiments.

### Dafadine-A promotion of dauer development

Dafadine-A (Sigma-Aldrich) was prepared as a 10 mM stock in DMSO and added to molten NGM, to a final concentration of 10 µM dafadine A (0.1 % DMSO v/v). Control plates were supplemented with the same final volume of DMSO (0.1 % v/v). All plates were seeded with 50 µl of OP50-1 and incubated overnight at room temperature. At low population density, *A. freiburgensis* hermaphrodite mothers produce female and male offspring (Kanzaki et al., 2017). Therefore ‘female-fated’ eggs were isolated from uncrowded plates (as described above) and added to the bacterial lawn of NGM plates, supplemented with dafadine-A or DMSO alone (4 plates for each condition with approximately 30 eggs per plate). Plates were incubated at 20 °C for 48 h, before the sex of the progeny was determined by phenotype. Females were physically larger and had reached adulthood, whilst those converted to hermaphrodite remained in the dauer larval stage. All non-male progeny was subsequently moved to NGM plates (without dafadine-A or DMSO) to continue development and examined daily for intestinal phenotype. In total, 390 individuals were scored (201 with dafadine-A, 189 control) from 3 individual experiments.

### Gene expression analysis

#### De novo transcriptome assembly

##### Sample preparation

Three biological replicates were generated for each RNA-seq condition (unmated females and hermaphrodites). Females and hermaphrodites were identified as described above. Approximately 150 individuals at the day 2 stage of adulthood were collected per sample into 1.5 ml tubes containing 200 µl of M9 buffer (Stiernagle, 2006). The nematodes were washed 3 times in M9 buffer. After the final wash, the supernatant was removed, 200 µl of Trizol® (Ambion) was added, and the sample stored at -80 °C. RNA was extracted as described previously (Adams et al., 2019). Residual genomic DNA was removed using Turbo DNase (Thermo Fisher Scientific) and samples cleaned with the RNA Clean and Concentrator kit (Zymo Research) according to the manufacturer’s instructions. Libraries were prepared with the TruSeq RNA Library Prep Kit v2 (Illumina) by the Genomics Facility at Warwick University.

##### RNA sequencing and Transcriptome Assembly

RNA-seq was performed on the Illumina HiSeq 4000 platform, generating a mean of 24.3 million 150 base pair end reads per replicate (Wellcome Trust, Oxford, UK). General assessment of the RNA-seq libraries was performed using FastQC (Andrews, 2010) and the raw reads from each library were pre-processed using Trimmomatic (version 0.36, TruSeq3-PE-2 adapters and the following parameters (“HEADCROP:15 SLIDINGWINDOW:5:20 MINLEN:36”) (Bolger et al., 2014). *De novo* transcriptome assembly was conducted with Trinity (v 2.8.3) including the jaccard-clip parameter (Grabherr et al., 2011; Haas et al., 2013). Potential contamination was removed using the DeconSeq program (Schmieder and Edwards, 2011) and CD-HIT-EST used to reduce redundancy in the assembly using a sequence identity threshold of 0.95 (Huang et al., 2010).

### Differential Gene Expression and GO term Analysis

Transcript abundance quantification was estimated within the Trinity software package (Haas et al., 2013) using the genome-free alignment-based quantification method RSEM (Li and Dewey, 2011). Processed reads were aligned to the *de novo* transcriptome with Bowtie2 and transcript abundance was estimated with RSEM, all within the Trinity package (v2.8.3). Differential gene expression analysis was conducted with EdgeR (also within Trinity v2.8.3). An adjusted p-value of 0.01 and an absolute log2 fold change (FC) of 1 were used to define differentially expressed genes.

The Trinity (v2.6.6) “gene to trans map” script was used to generate the gene to transcript map (Haas et al., 2013). Blast+ (v2.5.0), TransDecoder (v3.0.0) and hmmer (v3.1b2) were used to generate the relevant databases for Trinotate (v3.1.1) which was used to produce the annotations (Bendtsen et al., 2004; Bryant et al., 2017; Finn et al., 2011; Haas et al., 2013). We carried out an enrichment analysis (Fisher exact test) using the R (v3.5.1) package topGo (v3.8) on each of the lists of differentially expressed genes. This identified GO terms that are overrepresented (p < 0.01) relative to the entire transcriptome.

### Normalisation gene identification and qRT-PCR Analysis

Optimal normalisation genes were identified from the RNA-seq data using the NormFinder software (Andersen et al., 2004). This algorithm ranks candidate normalization genes according to their expression stability in all samples in a given experiment. *Afr-myosin* (TR7316|c0_g1_i1) and *Afr-tubulin* (TR12573|c1_g1_i1) were identified as the two top candidates. *Afr-tubulin* encodes for a tubulin-like protein (shares 98% identity with Cel-BEN-1) and *Afr-myosin* encodes for a myosin-like protein (shares 71% identity with Cel-HUM-5).

#### qRT-PCR Analysis

Samples were collected and RNA extraction conducted as described above. RNA was treated with DNase I (Sigma) to remove residual genomic DNA. cDNA synthesis was performed with 0.5 µg of RNA using random primers (Promega) and the MMLV reverse transcriptase enzyme (Promega) following the manufacturer’s instructions. qRT-PCR was conducted using the Stratagene Mx3005P detection system (Agilent Technologies) and GoTaq qPCR mix (Promega). Expression levels were calculated relative to the normalisation genes *Afr-myosin* or *Afr-tubulin*. All primer sequences used are listed in Supplemental Table S1.

## RESULTS

### *A. freiburgensis* unmated females exhibit mate-searching behaviour that is suppressed in reproducing females and hermaphrodites

If females and hermaphrodites are specialised in their roles, we would expect to see difference in their behaviours. Females, due to sexual morph specialisation (Charlesworth, 1999), would spend more time looking for a male. We tested this directly with behavioural assays measuring chemotaxis to male pheromones and food leaving behaviour.

There was a significant interaction between reproductive status and age on whether animals were attracted to male pheromones (logistic regression: χ^2^ = 10.113, d.f. = 4, p = 0.0386) (Figure 2A). Unmated females exhibited a high level of attraction to male pheromones at all time points, while actively reproducing hermaphrodites showed no attraction. Mated females became increasingly attracted to the male pheromone as they became sperm depleted (approximately 48 h after mating) and began to lay unfertilised oocytes (z= -2.390, p = 0.0444). By day 3, the chemotaxis index of mated females was comparable to that of virgin females (z= 1.748, p = 0.1875).

**Figure 2.**
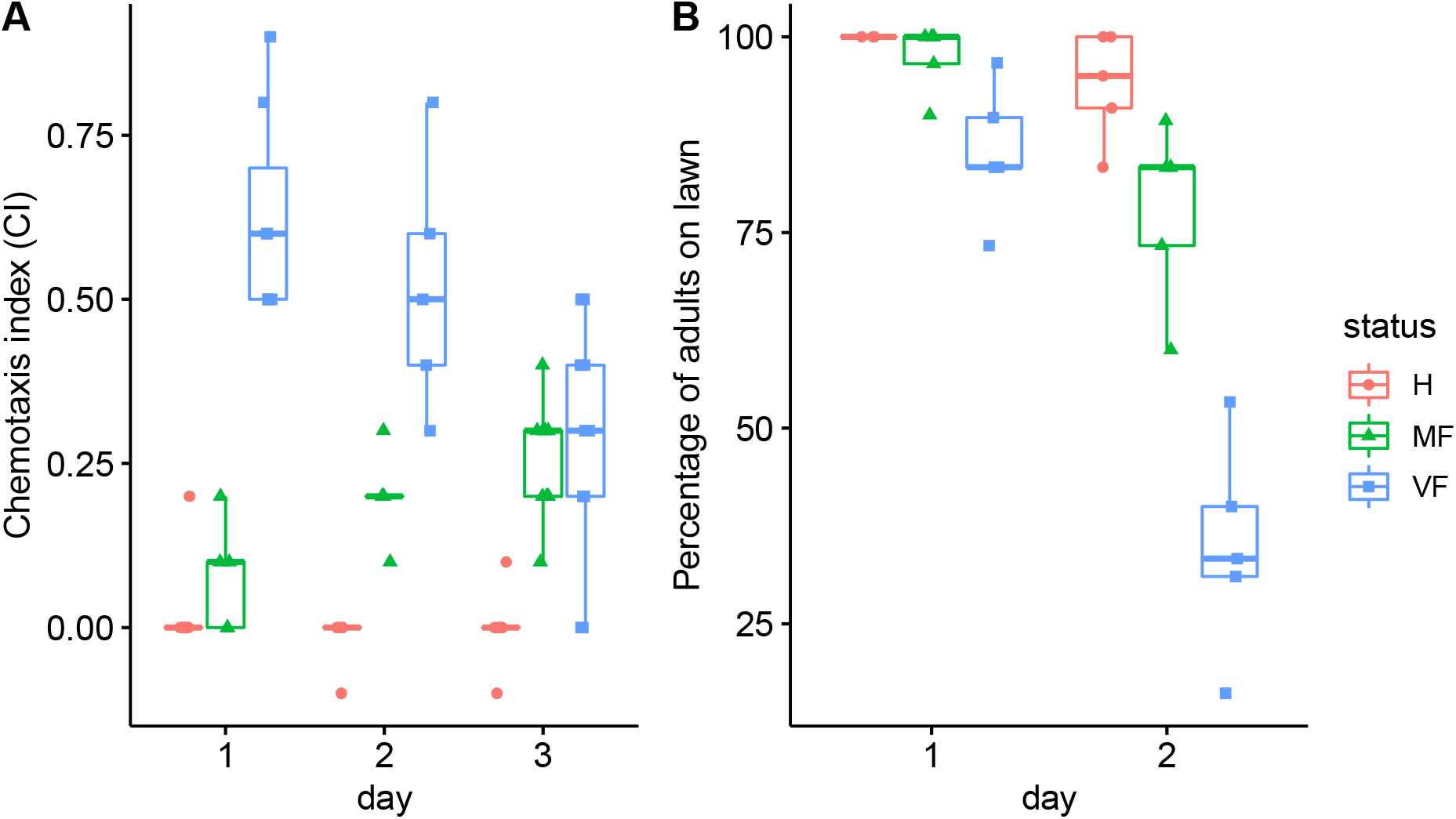
The *A. freiburgensis* female mate-searching behaviours change according to its reproductive status. (A) The chemotaxis index assay was used to measure attraction of hermaphrodites (H), mated females (MF) and virgin females (VF) to male supernatant (pheromone) at different days of adulthood. (B) Leaving behaviour out of the bacterial lawn varies according to reproductive status, sexual morph and days of adulthood.

Nematodes are usually cultured in a Petri dish containing agar seeded in the centre with a patch of bacterial lawn used as a food source. Food-leaving behaviour, whereby individuals leave the central food source to search surrounding areas of the culture plate where no food is present, is sometimes associated with the search of the nematode for a mating partner (Lipton et al., 2004). We found a significant interaction between reproductive status and sexual morph type on food-leaving behaviour (z = 27.751, d.f. =2, p < 0.0001) (Figure 2B). 48 h after the start of the experiment unmated females showed high levels of food-leaving behaviour compared with hermaphrodites (z = 11.639, p < 0.0001) and mated females (z = 7.125, p < 0.0001). Mated females also showed increased food-leaving behaviour compared to hermaphrodites after 48 hours (z = -3.575, p = 0.001). In contrast, no increase in leaving was observed in hermaphrodites over time (z = -1.338, p = 0.1810), suggesting that female leaving was not driven by food depletion.

### *A. freiburgensis* females compensate for the high investment in mate-finding by limiting intestinal development until mated

We hypothesised that females compensate for investing in costly mate-searching, by redirecting resources from other processes. In nematodes, the intestine is the major metabolic organ (McGhee, 2007). Consistent with reduced investment in metabolism, *A. freiburgensis* virgin female intestines are less developed compared to hermaphrodites at the same developmental stage (Figure 3).

**Figure 3.**
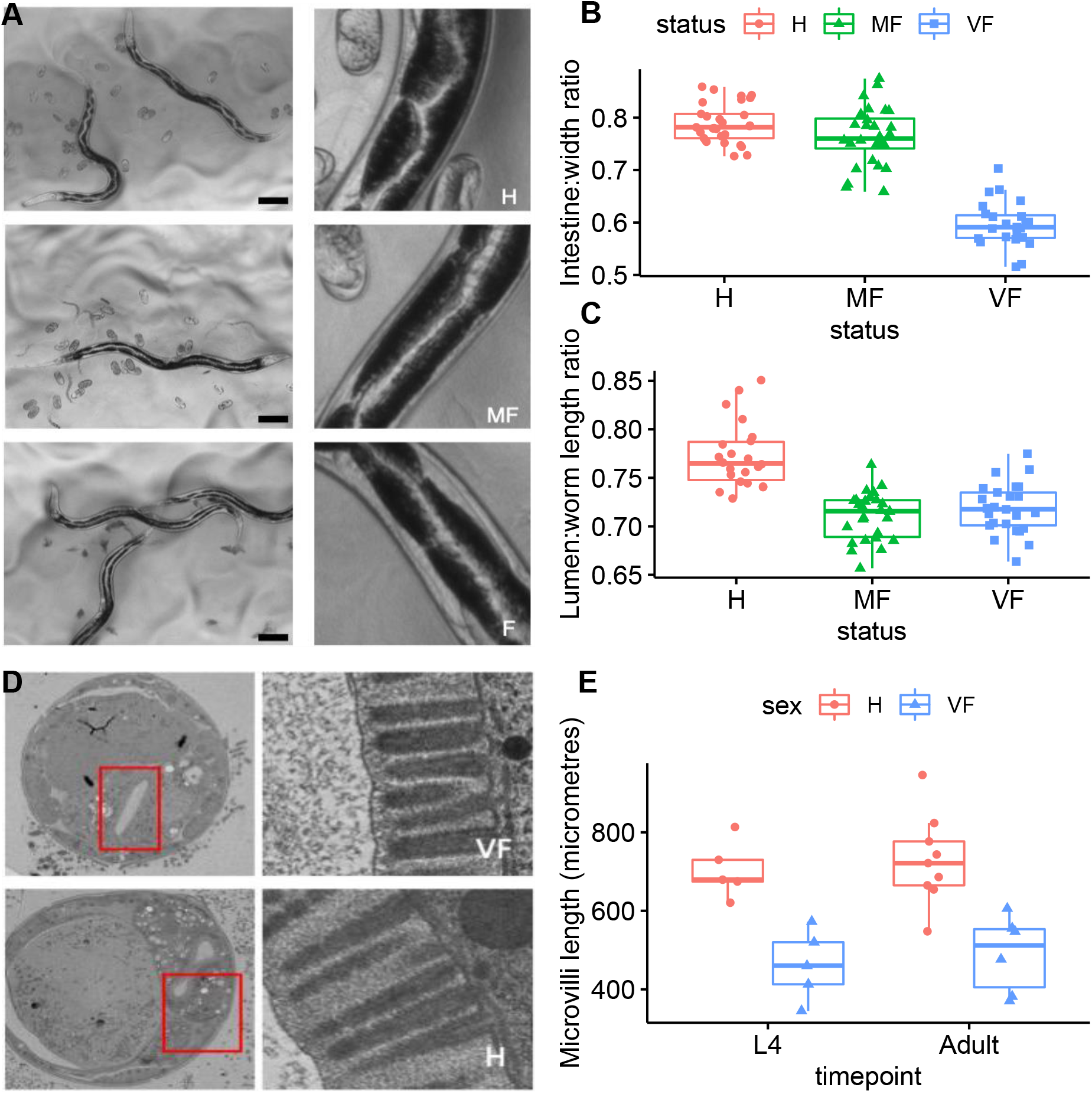
*A. freiburgensis* females limit investment in intestinal development until actively reproducing. (A) Representative examples of intestinal phenotype in H (top row), MF (middle row) and VF (bottom row) at day 2 of adulthood. The intestine is highlighted by the pairs of highly pigmented intestinal cells surrounding the intestinal lumen. Scale bar 100 μM. Measurements of intestinal width (B) and intestinal lumen length (C) on day 2 of adulthood. (B) Intestinal width was measured at the third intestinal cell from the tail and normalised to the whole nematode width at the same location. (C) Lumen length was measured from the pharynx bulb to anus and normalised to nematode length. (D) Transverse cross section EM images of VF and H (L4 larval stage shown). The microvilli brush border and glycocalyx layer is thicker in the H (right panel). The rectangle highlights the lumen cavity. (E) Intestinal microvilli length in VF and H, at the final larval stage before adulthood (L4) (see Figure 1) and day 0 of adulthood. H (hermaphrodite), MF (mated female), VF (virgin female).

Sexual morph had a significant effect on thickness of intestine (F_2,77_ =126.6, p=2 × 10^−16^) (Figure 3 B), length of lumen (F_2,70_ =33.86, p=5.18 × 10^−11^) (Figure 3 C) and length of microvilli (F_1,21_ =36.539, p=5.34 x10^−6^) (Figure 3 E). Compared to the hermaphrodite, the intestine of the virgin female is significantly thinner (Figure 3 B) (H - VF: t = 14.908, p < 0.0001). It also has a straighter and shorter lumen (Figure 3 C) (H - VF: t = 6.666, p < 0.0001), with shorter microvilli (Figure 3 D, E). Once mated, the female’s intestine expands (Figure 3 B) (MF – VF: t = 12.885, p < 0.0001), although the lumen length does not significantly increase (Figure 3 C) (MF – VF: t = -0.954, p = 0.608). The elongated lumen of the hermaphrodite is formed by an alternate arrangement of pairs of highly pigmented pyramidal intestinal cells that surround the lumen cavity that cause the lumen to twist and turn (Figure 3A). This organisation produces the characteristic zigzag pattern of the hermaphrodite intestine (Figure 3A).

### The enlarged and specialised intestine of the hermaphrodite promotes increased nutrient absorption

We hypothesised that the enlarged and specialised intestine of the hermaphrodite (and to a lesser extent the mated female) promotes increased nutrient absorption to meet the high energy demands of reproduction in *A. freiburgensis*. In *C. elegans*, excess energy is predominantly stored in neutral lipid vesicles in the intestine and epidermis (Mullaney and Ashrafi, 2009). We predicted that in *A. freiburgensis*, if the hermaphrodite intestine was better suited to nutrient uptake, hermaphrodites would have higher lipid stores than mated females in well-fed conditions.

Although staining of live nematodes with the neutral lipid dye Nile red is problematic, due to the sequestering of ingested dye into lysosome-related organelles (LROs) (O’Rourke et al., 2009), Nile red staining of fixed samples faithfully highlights neutral lipid stores in *C. elegans* (Mak, 2012) and *P. pacificus* (Kroetz et al., 2012). Therefore, we employed the fixed Nile red staining method (Pino et al., 2013) to study neutral lipid stores in unmated females, mated females and hermaphrodites. Age and mating status interactively had a significant effect on neutral lipid storage in *A. freiburgensis* (F_3,138_ =1.24 × 10^12^, p < 2 × 10^−16^) (Figure 4). Females entered adulthood (day 0) with high lipid stores, which were maintained if unmated (0 – 2 days: t = 0.901, p = 0.6406) but significantly decreased once they started to produce offspring (day 1 MF – VF: t = -4.227, p = 0.001). As females are less likely to leave the food source once mated (Figure 2B), depletion of energy stores is likely to be due to the redirection of resources and not due to reduced nutrient uptake. In contrast, hermaphrodite lipid store levels were low as they exited the non-eating dauer stage and entered adulthood (day 0) but rose steadily to significantly exceed those of unmated females by mid-adulthood (day 2) (H – VF : t = 5.222, p < 0.001). The hermaphrodite intestine may promote more efficient nutrient uptake as the elongated lumen and longer microvilli could increase the surface area for absorption. Not only would this allow the hermaphrodite to replenish energy stores depleted during diapause, but it could facilitate the production of progeny in less nutrient-rich surroundings.

**Figure 4.**
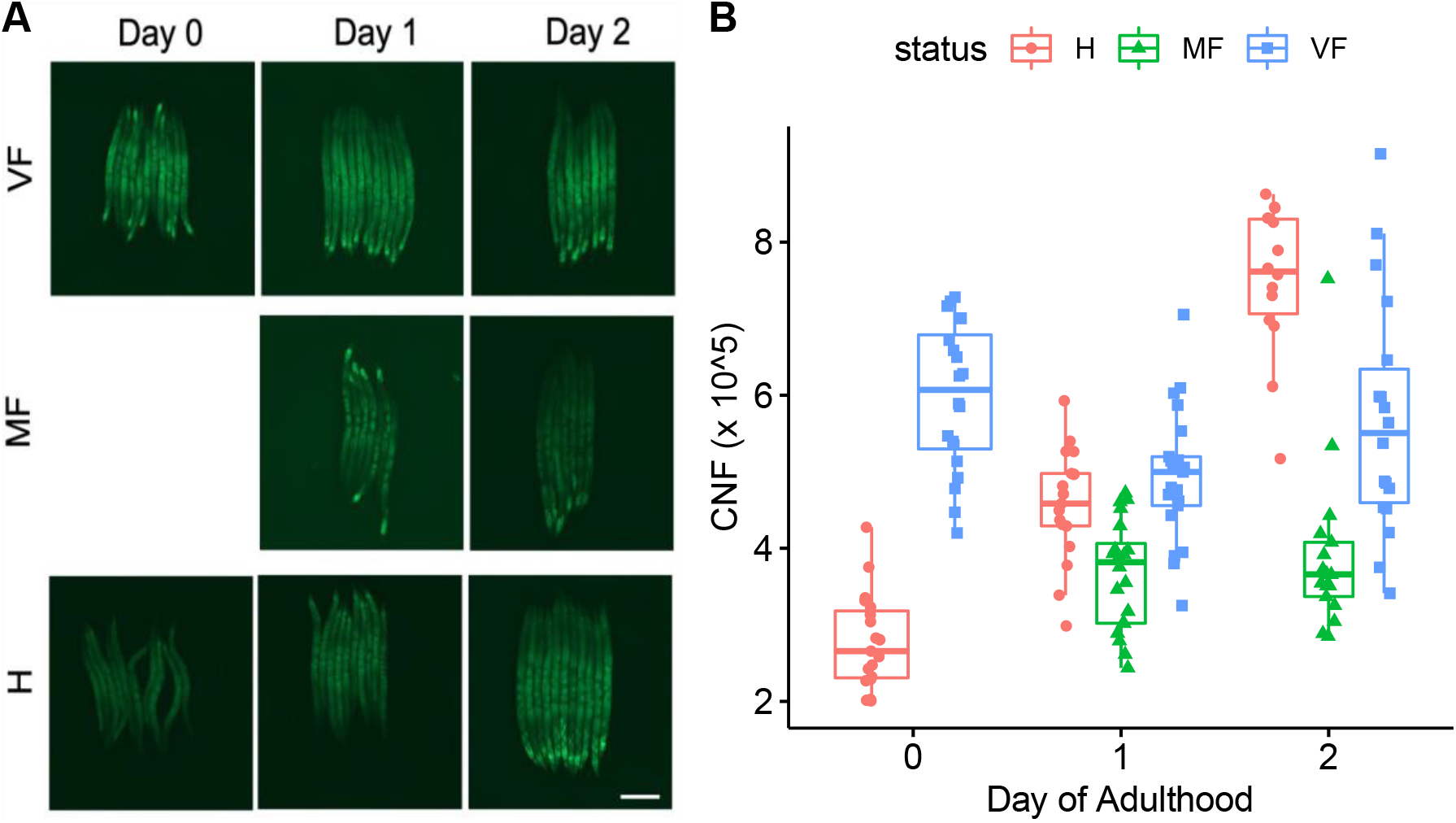
Neutral lipid stores fall in mated females, but rise in hermaphrodites, during peak egg production in *A. freiburgensis*. (A) Representative examples of fixed Nile red staining of lipid stores in VF, MF and H on day 0,1 and 2 of adulthood. MF were mated overnight (16 h) from L4 larval stage to day 0 of adulthood, in a ratio of 5 females to 1 male. Images taken with a 1 s exposure time. (B) Quantification of lipid staining. Corrected nematode fluorescence (CTNF) was calculated for individuals from 3 independent experiments (see methods for details). H (hermaphrodite), MF (mated female), VF (virgin female). Scale bar 100 μM.

### Passage through dauer links mating system and optimised intestinal morphology in *A. freiburgensis*

At the L3d/L4 stage of larval development *A. freiburgensis* hermaphrodites, the zigzag intestinal pattern resulting from the positioning of the pyramidal cells becomes apparent (Figure 5A). In contrast, the L3 female intestinal cells are cuboid and the lumen straight. These differences in intestinal morphology continue into adulthood. As *A. freiburgensis* hermaphrodites pass through an obligate dauer, a larval stage where the *C. elegans* intestine undergoes remodelling (McGhee, 2007), we postulated that passage or bypassing of the dauer larval stage may regulate sexual morph-specific intestinal development. Therefore, we exploited the ability to chemically induce or inhibit dauer development to determine if passage through dauer was sufficient to modulate intestinal development (see methods for details).

**Figure 5.**
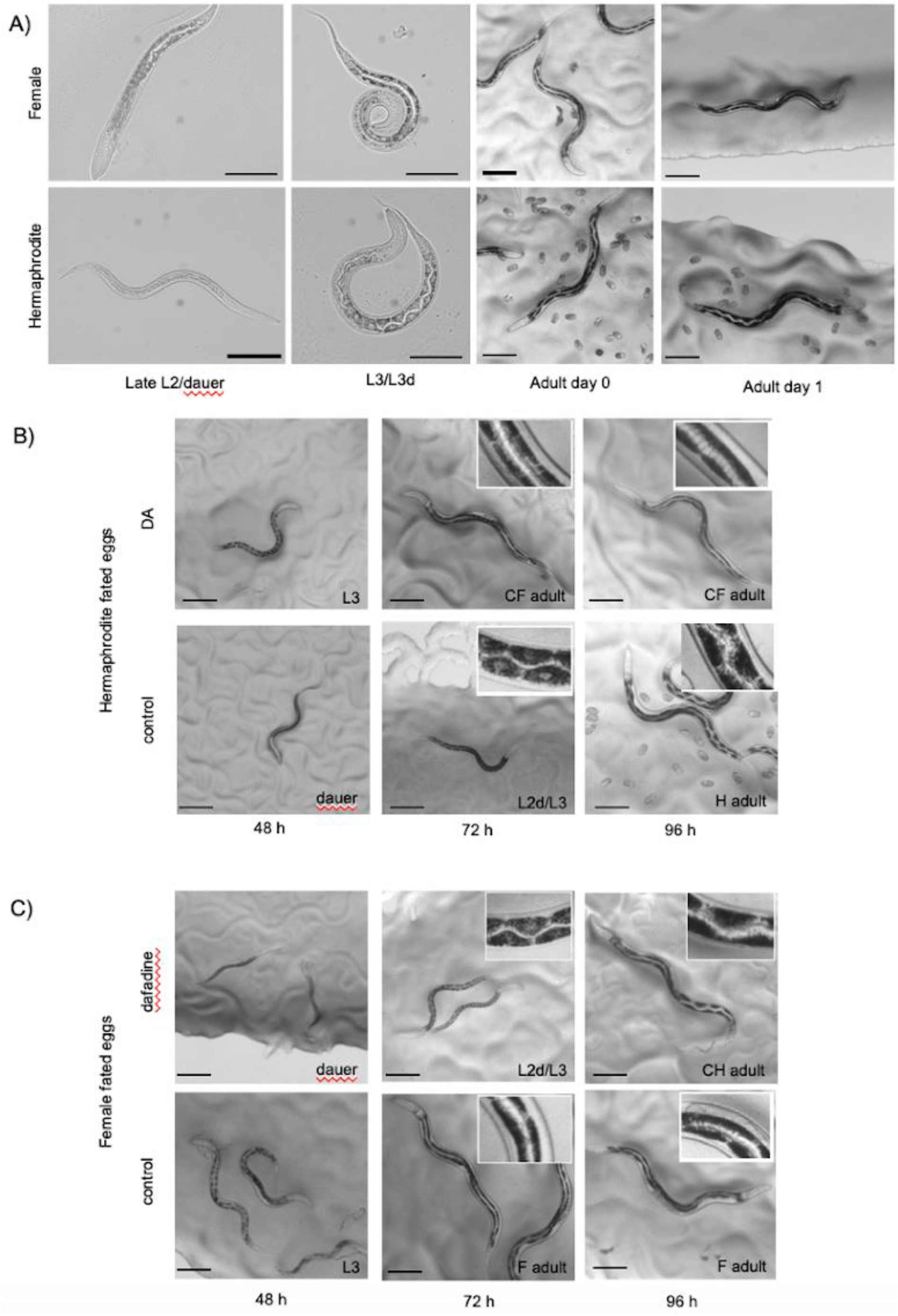
Passage through dauer larval development is sufficient to induce the distinctive hermaphrodite intestinal phenotype in *A. freiburgensis*. (A) Female and hermaphrodite intestinal development diverged at the L2/dauer stage. The chemical inhibition (B) or induction (C) of dauer development illustrates that the hermaphrodite intestinal phenotype in *A. freiburgensis* is intrinsically linked with passage through dauer. (B) Hermaphrodite-fated eggs, incubated on NGM supplemented with DA (10 μM in 1% ethanol), failed to enter dauer and were diverted to the female fate (top panel). These converted females (CF) exhibited the wild-type female intestinal phenotype. Hermaphrodite-fated eggs incubated on the control plates (1% ethanol alone) entered dauer and developed the characteristic hermaphrodite intestinal phenotype (bottom panel). (C) Female-fated eggs exposed to dafadine-A (10 µM (0.1 % DMSO v/v)) were forced into dauer development and diverted to hermaphrodite development (top panel). The converted hermaphrodites (CH) exhibited the zigzag pattern and pyramidal intestinal cell shape of wild-type hermaphrodites. On control plates, (0.1 % DMSO v/v), female-fated eggs followed normal female reproductive and intestinal development. Scale bars all 100 μM.

Previously, we have shown that addition of the hormone Δ7-dafachronic acid (DA) is sufficient to block dauer development in *Auanemna rhodensis* (Chaudhuri et al., 2011). To test if the dauer pathway can be manipulated in *A. freiburgensis*, hermaphrodite-fated eggs were collected from crowded plates and incubated either with DA (10 μM in 1% ethanol (v/v)) or on control plates (1% ethanol alone (v/v)). With supplemented DA, no dauers were observed (n=409), whilst on control plates 86 % non-males (n=221) passed through dauer.

To determine if addition of the DAF-9 inhibitor dafadine-A (Luciani et al., 2011) is able to promote the dauer fate in *A. freiburgensis*, female-fated eggs were collected from isolated hermaphrodite mothers and incubated, either on NGM plates with dafadine-A (10 μM, 0.1% DMSO (v/v)) or on control plates (0.1 % DMSO alone (v/v)). In the presence of dafadine-A, all non-male progeny passed through dauer (n= 201). No dauers were observed on the control plates (n=189). Consistent with previous studies, all individuals that passed through dauer developed into hermaphrodites, whilst non-males that bypassed dauer became female adults ((Kanzaki et al., 2017).

Passage through dauer was sufficient to induce the hermaphrodite intestinal phenotype, regardless of sexual morph at birth (Figure 5B-C). Female-fated individuals that were chemically forced through dauer exhibited the pyramidal cell shape (Figure 5B). In contrast, if dauer development was blocked in hermaphrodite-fated larvae, they exhibited the reduced intestine, straight lumen and intestinal cell shape of female-born larvae (Figure 5C).

### A. freiburgensis females and hermaphrodites exhibit differential gene expression in pathways related to mate-finding and metabolism

We carried out RNA-seq analysis to determine which biological pathways are differentially regulated between females and hermaphrodites. As we predicted that the major cost for females will be reproductive assurance, we compared unmated females with hermaphrodites at the same stage of adulthood (2 days of adulthood).

We identified 819 differentially expressed genes (with a cut-off of 2-fold change and p <0.01), with 275 upregulated in females (Supplemental Table S2) and 544 upregulated in hermaphrodites (Supplemental Table S3). Upregulated transcripts in hermaphrodites showed gene ontology (GO) term enrichment for genes associated with reproduction, embryo/larval development and ovulation (Supplemental Figure S1). There was also enrichment for GO terms associated with metabolic processes (including lipid catabolism and proteolysis) and digestion. In addition, hermaphrodites showed enhanced expression of genes associated with cuticle development and defence responses. Many upregulated transcripts in the female were associated with signalling processes and neuronal regulation (Supplemental Figure S2). Many of these genes have been directly implicated in mating and mate-searching behaviour (Table 1).

**Table 1.**
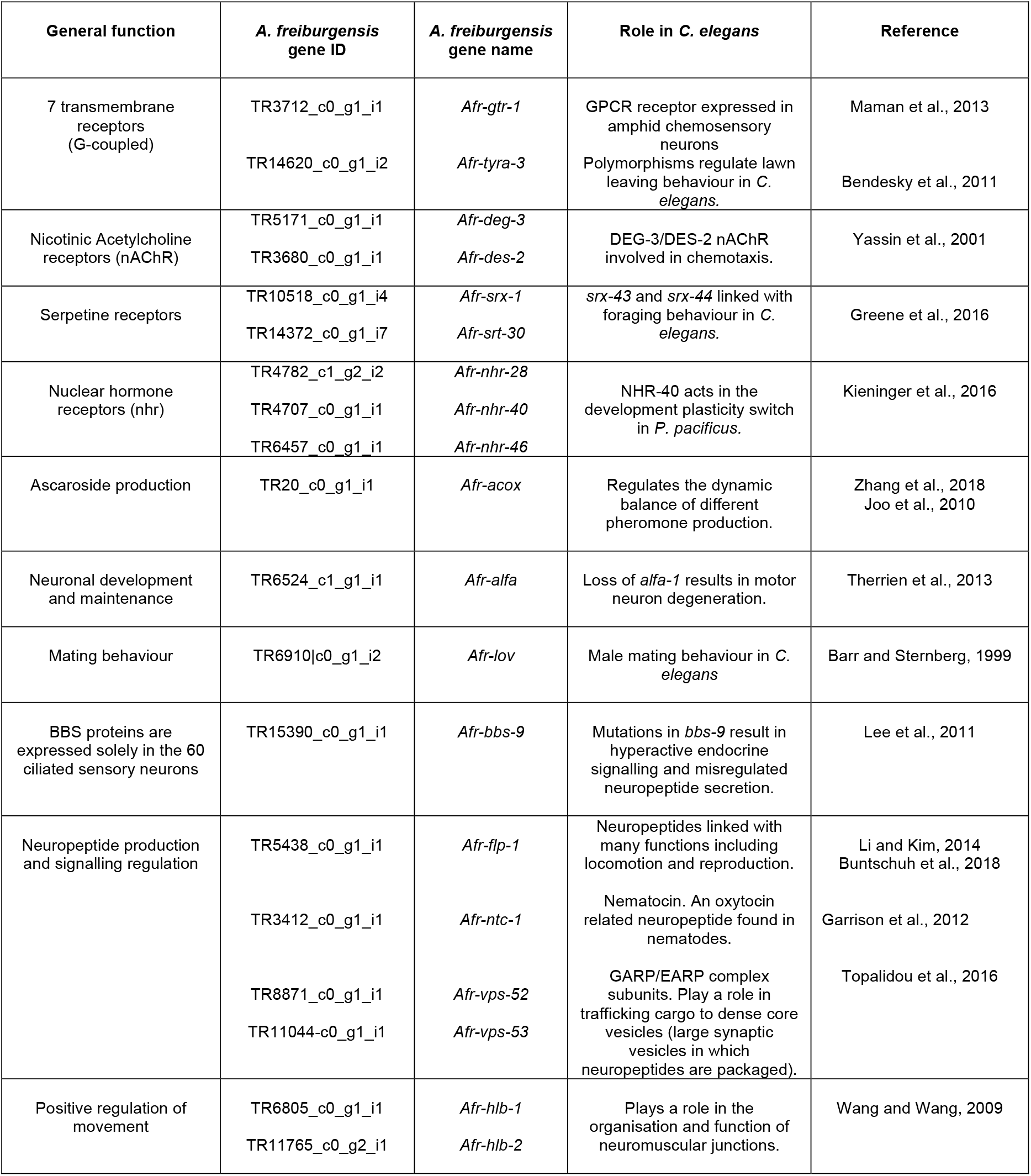
Neuronal and signalling genes upregulated in virgin females compared to hermaphrodites in *A. freiburgensis.*

### Signalling pathway genes upregulated in virgin females compared to hermaphrodites were downregulated after mating

Our data suggest that unmated females upregulate specific neuronal development and signalling pathways (See Table 1). If these pathways are a specific female adaptation to drive mate-searching behaviour, then mating should downregulate the expression of the identified neuronal/behavioural candidate genes. Therefore, we used qRT-PCR to analyse the expression of a subset of these candidate genes in virgin females, mated females and hermaphrodites, throughout peak egg production (Figure 6B-F, Supplemental Figure S3). All the neuronal-linked candidates tested (*Afr-ntc-1, Afr-tyra-3, Afr-deg-3, Afr-des-2* and *Afr-flp-1*) were suppressed in females after mating, often to the level observed in hermaphrodites. Interestingly, from day 2 to 3 of adulthood, the expression levels of many of the neuronal candidates increased in mated females, which coincided with the majority of these individuals ceasing egg laying, due to depletion of sperm stores. This suggests that this suite of genes can be reversibly repressed by reproduction. In contrast, *Afr-ges-1*, a homologue of the gut-specific esterase *ges-1* (Edgar and McGhee, 1986) is suppressed in females until reproduction starts (Figure 6A).

**Figure 6.**
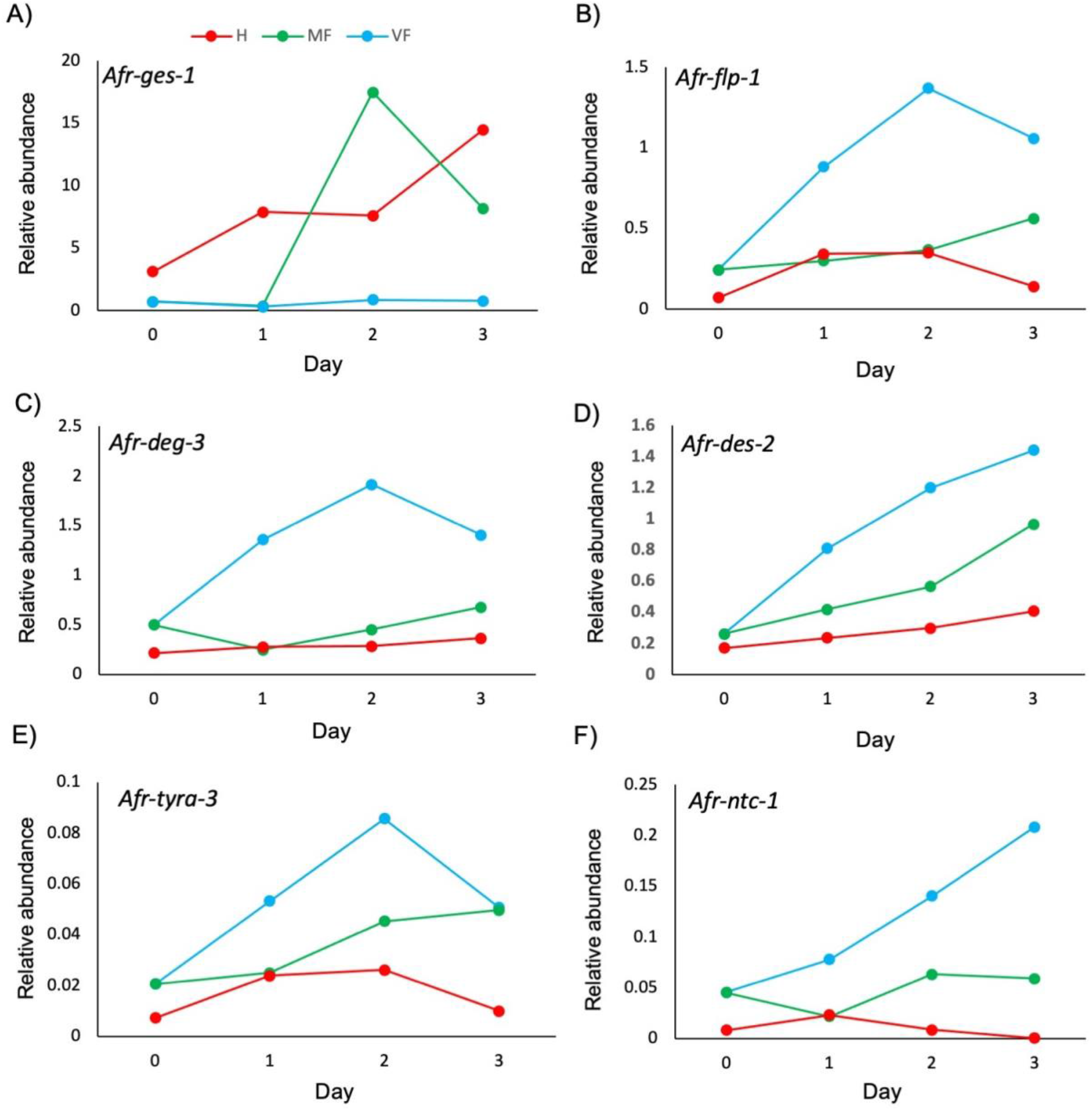
Non-reproducing females exhibit increased expression of mate-searching candidates. Representative example of expression time course analysis of *Afr-ges-1* (A) and candidates upregulated in virgin females *Afr-flp-1* (B), *Afr-deg-3* (C), *Afr-des-2* (D), *Afr-tyra-3* (E) and *Afr-ntc-1* (F). Transcript levels were determined by quantitative reverse transcription PCR (RT-PCR) and expressed relative to the normalisation gene *Afr-myosin*. Day is day of adulthood. VF (virgin females), MF (mated females), H (hermaphrodite). A replicate time course is shown in Supplemental Figure S3.

## DISCUSSION

We hypothesised that the female (obligate outcrossing) and hermaphrodite (predominantly selfing) morphs of *A. freiburgensis* play distinct roles and that each exhibits adaptations to minimise the cost of their specific life history, allowing the stable maintenance of trioecy. Unmated females specialise in finding males, showing the upregulation of genes implicated in mate-finding, increased attraction to male pheromones and enhanced mate-searching behaviour. Females may compensate for this high investment in mate finding by limiting metabolism and intestinal development till they are mated.

Hermaphrodites, on the other hand, have specialised in reproduction and energy acquisition. Passage through a migratory larval state is sufficient to ensure that a hermaphrodite enters adulthood able to self-fertilise with several intestinal adaptations likely to promote nutrient uptake. The hermaphrodite’s enlarged and specialised intestine could allow the individual to not only replenish energy stores depleted during migration, but also meet the high energy demands of reproduction. Consistent with hermaphrodite specialisation in energy acquisition they also show upregulation of genes involved in metabolic processes.

*A. freiburgensis* unmated females show upregulation of genes whose homologs are associated with chemoattraction, pheromone production, lawn-leaving and other mating behaviours (Barr and Sternberg, 1999; Bendesky et al., 2011; Garrison et al., 2012; Joo et al., 2010; Yassin et al., 2001; Zhang et al., 2018). Suppression of these genes in mated females and hermaphrodites correlated with reduced lawn-leaving and chemoattraction behaviour in *A. freiburgensis*. Lawn-leaving and attraction to male pheromones have been described in other obligate outcrossing nematodes, suggesting they represent conserved mate-searching traits (Borne et al., 2017; Chaudhuri et al., 2015; Choe et al., 2012; Duggal, 1978; Lipton et al., 2004). As with *A. freiburgensis*, mating suppresses many of these drives, including female lawn-leaving and attraction to male pheromones, indicating that these female behaviours are governed by reproductive status and are associated with the need to mate (Borne et al., 2017; Chaudhuri et al., 2015; Duggal, 1978; Lipton et al., 2004).

In most organisms, mate-searching behaviour is predominantly associated with males, as females are thought to rarely require more than one mating to fertilise all their eggs (Andersson, 1994). However, models predict that females could benefit from mate searching if females are at risk of not mating or would benefit from multiple matings (Kvarnemo and Simmons, 2013; Parker and Birkhead, 2013; Rhainds, 2010). As males are under-represented in *A. freiburgensis* populations and time-limited mated females became sperm depleted, natural selection could favour females who actively search for males.

Investment in mate-searching behaviour is costly. Our data suggests that unmated *A. freiburgensis* females may meet this cost by reallocation of resources from metabolism and intestinal expansion. Although adult tissue is often considered to be homeostatically maintained at a constant size, there are a growing number of examples of post-developmental organ plasticity to meet the energy demands of reproduction. Expansion of the alimentary tract during pregnancy and/or lactation has been observed in diverse mammalian species (Hammond, 1997; Speakman, 2008), and hormones released post-mating in *Drosophila* promote an increase in the midgut (an organ which plays a similar role to the mammalian small intestine), enhancing reproductive output (Reiff et al., 2015). Interestingly, although the female intestine expands post-mating in *A. freiburgensis*, it does not develop all the characteristics of the hermaphrodite intestine triggered by passage through dauer. Delayed or reduced intestinal development may explain why female lipid stores were depleted, whilst hermaphrodite levels increased, during peak egg reproduction. As *A. freiburgensis* females are predominantly produced in uncrowded conditions (Kanzaki et al., 2017) when food supplies are likely to be plentiful, reducing intestinal development, and by association nutrient uptake, may be an ‘acceptable’ risk.

In conclusion, *A. freiburgensis* exhibits developmental and behavioural plasticity dependent upon both sexual morph and mating status. In simple terms, females are not merely hermaphrodites who are unable to produce sperm, or vice versa. Although we cannot discount that trioecy in *A. freiburgensis* is an evolutionary transient state, it is feasible that selection for specific female and hermaphrodite adaptations drives the stable co-existence of both mating strategies. Trioecy has been identified in a growing number of animal species including; *Hydra viridissima* (Kaliszewicz, 2011), the marine mussel, *Semimytilus algosus* (Oyarzún et al., 2020) and the sea anemone *Aiptasia diaphana* (Armoza-Zvuloni et al., 2014). The study of sexual morph specialisation in this growing number of trioecious animal species will allow the development of more robust mathematical models to investigate the long-debated evolutionary enigma of mixed mating strategies.

## ACKNOWLEDGMENTS

The authors would like to acknowledge the help of the Media Preparation Facility in the School of Life Sciences at the University of Warwick. A.P.-d.S. and S.A. were supported by a grant from BBSRC (BB/L019884/1). EBM was funded by NERC grant NE/N010019/1 and the Leverhulme Trust. ARCJ was supported by a BBSRC MIBTP DTP studentship.

## DATA ACCESSABILITY STATEMENT

Upon acceptance, RNA-seq data will be uploaded to the European Nucleotide Archive (ENA) (https://www.ebi.ac.uk/ena/browser/home).

## AUTHOR CONTRIBUTIONS

S.A. and A.P.S. designed the study. S.A. and P.P. conducted the experimental work except TEM. Data analysis was conducted by S.A., A.R.C.T and E.B.M. TEM experiments and analysis were conducted by M.K. The article was written by S.A. and E.B.M with input from all other authors.

## Notes

### Competing Interest Statement

The authors have declared no competing interest.

