## Supplemental_data for "Sexual morph specialisation in a trioecious nematode balances opposing selective forces"

**Supplemental Table S1** **Primer sequences used for qRT-PCR analysis**

| **Gene ID** | **Gene name** | **Primer name** | **DNA sequence (5’ to 3’)** |
| --- | --- | --- | --- |
| TR6856\|c0_g1_i1 | *Afr-ges-1* | UW376  UW377 | GCTCAACCACCTGTCGGAAACCT  CTTTGGGTGAGGAGTAGCGGCTG |
| TR5438_c0_g1_i1 | *Afr-flp-1* | UW567  UW568 | CATAGCAACTACGGCACAGGTAGC  TGCTTGTTCCATGGTCGACAGCA |
| TR5171_c0_g1_i1 | *Afr-deg-3* | UW571  UW572 | TGCTTCGACTGGCCTCATCAGTAC  TCCTGCAAAGCCAGCACTGTTTG |
| TR3680_c0_g1_i1 | *Afr-des-2* | UW569  UW570 | CGTTGGGTAGCCAAACTGGTTCG  CTTCTTGAGCCCTCCGCAGCTTC |
| TR14620_c0_g1_i2 | *Afr-tyra-3* | UW463  UW464 | ACGCGAACGAGATCAAGAGGGATG  TGGTACAATCCATGGAGGGGTGC |
| TR7316\|c0_g1_i1 | *Afr-myosin* | UW398  UW399 | AGGCCACTAGTCTCTGTAGCCCC  TCGATTGGTCTGCTGGGAACACG |
| TR12573\|c1_g1_i1 | *Afr-tubulin* | UW396  UW397 | TCGTTGACTTGGAACCCGGAACC  CCGGCTCCAGATTGTCCGAAGAC |

**Supplemental Table S2** **Genes upregulated in females**

(Excel sheet)

**Supplemental Table S3** **Genes upregulated in hermaphrodites**

(Excel sheet)

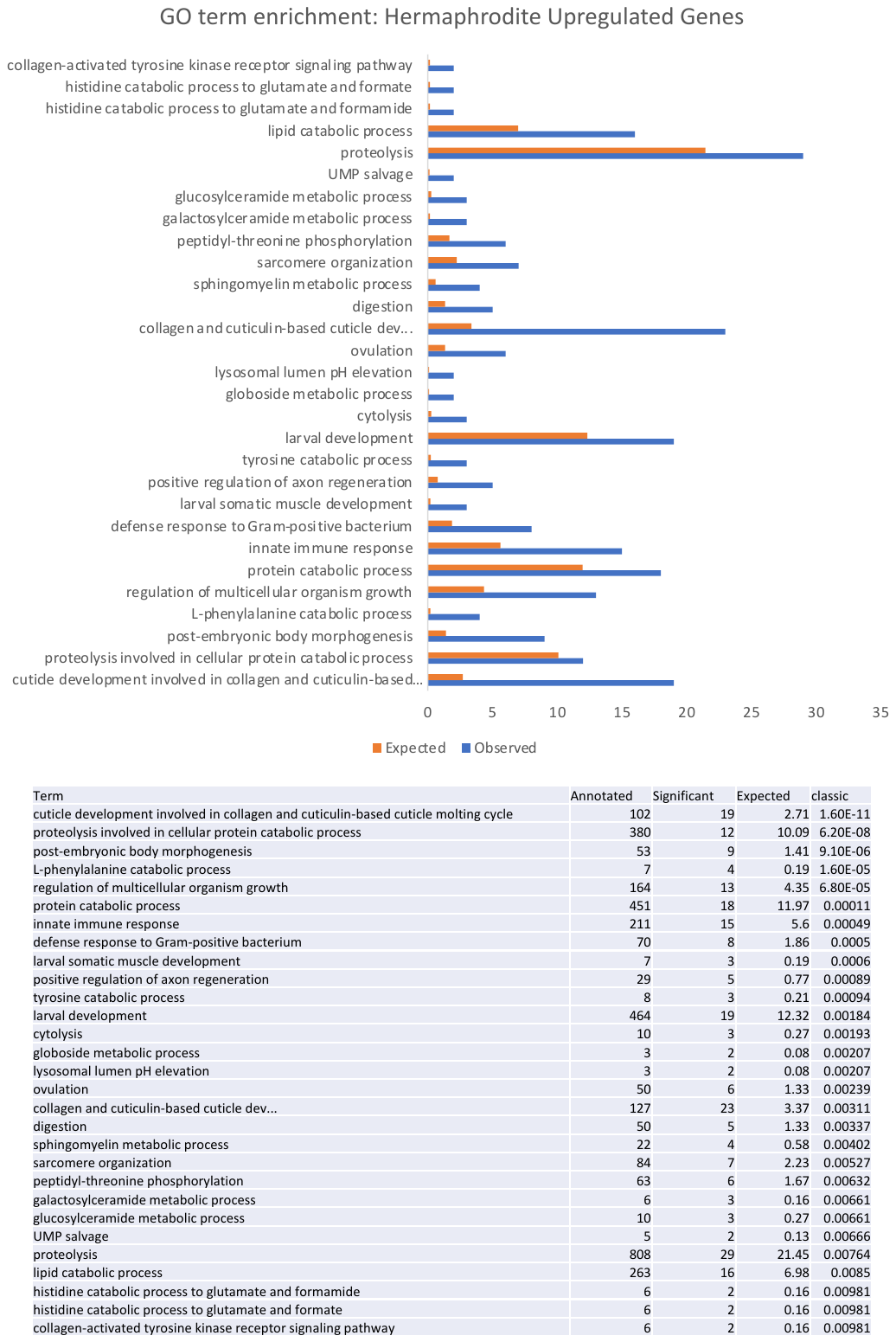
**Supplemental Figure 1.**

**GO term enrichment analysis of genes upregulated in *A. freiburgensis* hermaphrodites**

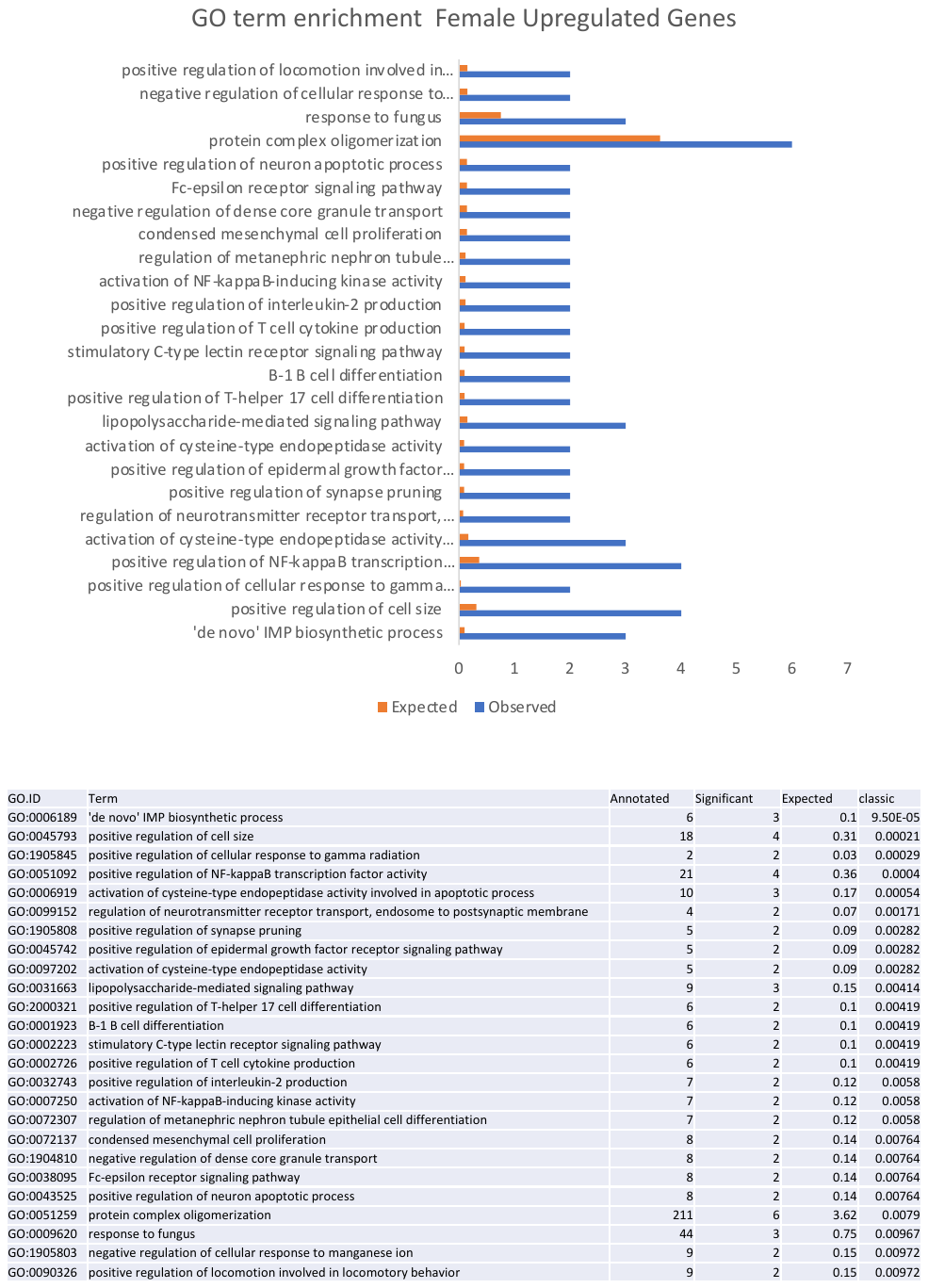

**Supplemental Figure 2.**

**GO term enrichment analysis of genes upregulated in *A. freiburgensis* females**

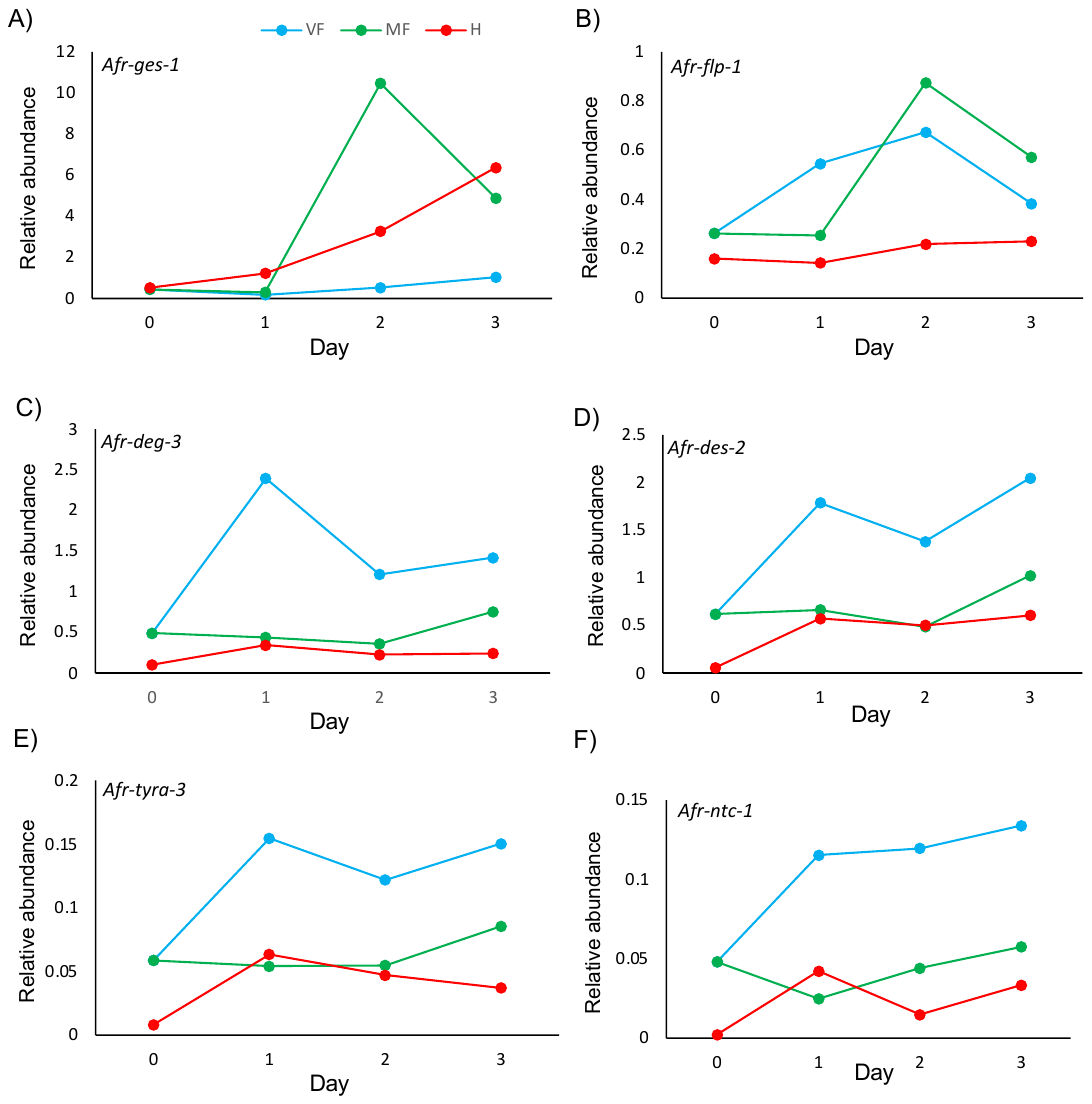

**Supplemental Figure 3. Non-reproducing females exhibit increased expression of mate-searching candidates** Representative example of expression time course analysis of *Afr-ges-1* (A) and candidates upregulated in virgin females *Afr-flp-1* (B), *Afr-deg-3* (C), *Afr-des-2* (D), *Afr-tyra-3* (E) and *Afr-ntc-1* (F). Transcript levels were determined by quantitative reverse transcription PCR (RT-PCR) and expressed relative to the normalisation gene *Afr-myosin*. Day is day of adulthood. VF (virgin females), MF (mated females), H (hermaphrodite).
